# Characterization of three *Yarrowia lipolytica* strains in respect to different cultivation temperatures and metabolite secretion

**DOI:** 10.1101/645242

**Authors:** S Hackenschmidt, F Bracharz, R Daniel, A Thürmer, S Bruder, J Kabisch

## Abstract

Despite the increasing relevance, ranging from academic research to industrial applications, only a limited number of nonconventional, oleaginous *Yarrowia lipolytica* strains are characterized in detail. Therefore, we analyzed three strains in regard to their metabolic and physiological properties and in respect to important characteristics of a production strains. A flow cytometry method was set up to evaluate their fitness in a rapid manner. By investigating different cultivation conditions and media compositions, similarities and differences between the distinct strain backgrounds could be derived. Especially sugar alcohol production, as well as a agglomeration of cells were found to be connected with growth at high temperatures. In addition, sugar alcohol production was independent of high substrate concentrations under these conditions. To investigate particular traits, including growth characteristics and metabolite concentrations, genomic analysis were performed. We found sequence variations for one third of the annotated proteins but no obvious link to all phenotypic features.

## Introduction

The growing awareness of human impact on the environment, especially the global effects of massive utilization of fossil resources, leads to a greater demand of sustainable substitutes. The usage of biomass for the production of renewable lipid derived products, such as hydrocarbons, could be a future-proof alternative. Although traditional oil crops are still the main source for renewable oil, microbes gain in importance, as they have shorter life cycle, do not dependent on arable land and do not necessarily compete with human or animal food production. Nonetheless, the high processing costs constitute a major barrier for an economically viable large-scale production of microbial oil (Braunwald *et al*. 2014; Parsons *et al*. 2018).

The selection of the microbial strain is one of the first steps to develop a competitive process. Therefore scientists recoursed to well studied organisms like *Escherichia coli* and *Saccharomyces cerevisiae* that are already widely applied in industrial processes. However, both organisms exhibit a rather low lipid content despite numerous metabolic engineering approaches (Ferreira *et al*. 2018; Teixeira *et al*. 2018; Xu *et al*. 2018). The exploration of nonconventional microorganisms which naturally accumulate lipids is likely a more promising strategy. These oleaginous microorganisms are taxonomically diverse and store lipids up to 80 % of their dry cell weight (Sitepu *et al*. 2013; Athenaki *et al*. 2018; Bellou *et al*. 2014). However, their industrial applicability needs to be proven and commonly genetic and molecular biology tools must be developed to transform them into modifiable production platforms (Alper and Stephanopoulos 2009; Abdel-Mawgoud *et al*. 2018).

The oleaginous yeast *Yarrowia lipolytica* is an established industrial host for the production of carotenoids and lipids and is currently developed into a microbial cell factory, as reviewed by Markham and Alper (Markham and Alper 2018). When e.g. nitrogen sources are depleted, oleaginous organisms like *Y. lipolytica* continue to assimilate carbon sources. As nitrogen is missing for protein and nucleic acid synthesis cell proliferation stops and the carbon is directed to synthesis of triacylglycerols which are stored in specialised compartments, the lipid bodies. Lipid production with different *Y. lipolytica* strains, its regulation and optimization is described in numerous publications (Bruder *et al*. 2018a; Lazar, Liu and Stephanopoulos 2018; Zeng *et al*. 2018).

In contrast to the increasing relevance of *Y. lipolytica* as a host, just a few well characterized strains are available or commonly utilized. Currently, a strong research focus lies on the natural isolates H222, W29, including its derivatives from the Po1 series and CBS6142-2 (Beopoulos *et al*. 2010; Bredeweg *et al*. 2017). High quality genomic sequences are likewise analyzed and provided just for a couple of strains (Devillers *et al*. 2016; Magnan *et al*. 2016; Pomraning and Baker 2015; Liu and Alper 2014; Dujon *et al*. 2004). Efforts to overcome this restricted view were recently made by (Egermeier *et al*. 2017). The authors could demonstrate a correlation between the isolation site and certain metabolic characteristics. *Yarrowia* strains isolated from dairy products specifically produced high amounts of polyols when grown under nitrogen limitations on glycerol media. To further expand the exploration of *Y. lipolytica* intraspecies diversity, we characterized and compared three different strains regarding physiological and metabolic properties. Therefore, two hitherto not characterized wild type isolates and the common laboratory strain H222 with a knockout of its ß-oxidation genes were chosen.

In detail, comparisons in terms of growth in different media and at different temperatures, as well as morphological differences at different growth stages were investigated. Additionally industrially relevant metabolites as well as lipids were quantified. Shotgun sequencing and analysis of reference mappings of all three examined strains was performed but did not yield any conclusive results.

## Materials and Methods

### Strains and cultivation conditions

Strains used in this work were *Y. lipolytica* SBUG-Y 63 and SBUG-Y 1889 (kindly provided by F. Schauer, Ernst-Moritz-Arndt University Greifswald, Germany) referred to as 63 and 1889, respectively. Additionally, the laboratory strain H222Δ*pox1-6* was used (Gatter *et al*. 2014).

Cultivations were performed with defined synthetic YNB media, consisting of 1.7 g L^−1^ yeast nitrogen base without amino acids (obtained from Sigma Aldrich) as well as lipogenesis inducible YSM medium (Bruder *et al*. 2018b) and complete medium YPD (Zimmermann 1975). YNB and YPD contained 2 % (w/v) and YSM 5 % (w/v) D-glucose. All pre-cultivations were performed in the corresponding media at 28 °C. Cultivations were performed at 28 °C or 35 °C and 180 rpm (51 mm amplitude) in a New Brunswick Innova 44 shaker. Measurements were stopped after 100 h of cultivation. Temperature settings were verified with a high precision thermometer (P4010, Dostmann Electronic GmbH). Growth rate analyzes were performed with RStudio version 1.0.153 using R version 3.5.1 (Ihaka and Gentleman 1996), tidyverse version 1.2.1 and grofit version 1.1.1-1 (using Gompertz model).

### Determination of extracellular metabolites

Metabolites of remaining cell free supernatant were analyzed by HPLC. Concentrations of D-glucose, citrate and polyols were determined by Perkin Elmer Series 200, using a Rezex^TM^ ROA-Organic Acid H+ (8 %) column (Phenomenex). References were purchased from Sigma Aldrich. The column was eluted with 5 mM sulfuric acid as mobile phase and a flow rate of 0.4 ml min^−1^ at 65 °C. Refractive index values were detected by RI-101 (Shodex). For data evaluation, TotalChrom Workstation/Navigator software (Perkin Elmer, version 6.3.2) was used. Data conversion and plotting was performed with RStudio as described above.

### Lipid analytics of cell extract

Samples of 1 ml volume were extracted by simultaneous disruption of cells using Retsch Mixer Mill MM 400 (20 min, frequence of 30 s^−1^ at RT) in 300 µL hexane/2-propanol (3:1, v/v) containing internal standard (5 mM tridecanoic acid) and 200 μl A. dest. After phase separation, direct transesterification was carried out by adding 500 μl 2 % (w/v) sulfuric acid in methanol and 50 μl 2,2-dimethoxypropane (water scavenger) followed by incubation at 60 °C for 2 h & 1400 rpm in a thermomixer (eppendorf Thermomixer comfort). Fatty acid methyl esters (FAME) were extracted with 300 µl hexane. The samples were analyzed with a Shimadzu Nexis GC 2030, on a Shimadzu SH-Rxi-5MS column (30 m, 0.25 mm, 0.25 µm) and detected by FID. The temperature of inlet and FID were set to 250 °C and 310 °C, respectively. The temperature program of the column oven consists of a five steps: (1) temp. 90 °C, hold 5 min; (2) rate 15, final temp. 190 °C; (3) rate 2.0, final temp. 200 °C, hold 1 min; (4) rate 0.5, final temp. 202.5 °C, hold 1 min; (5) rate 20, final temp. 300 °C, hold 5 min. For peak assignment, FAME mix from Sigma Aldrich (CRM18918) was used. For quantification, corresponding single FAMEs from Sigma Aldrich Fluka in the concentration range of 0.025 - 8 mM were measured. Data were processed using LabSolutions 5.92 and RStudio as described above.

### Flow Cytometry

Cytometric analysis was done using a Sony LE-SH800SZBCPL with a 488 nm argon laser. Photomultipliers for backscatter were set on 25.5 % with a forward scatter threshold of 0.20 % and a window extension of 50. The forward scatter diode was set on an amplification level of 6/12 and sample pressure was set so that events per second were kept under 30’000. Areas of scattering signals were brought to a near-normal form over the inverse hyperbolic sine transformation. Gating was avoided to ensure inclusion of agglomerates and hyphae. Advanced settings are shown in Figure S3.

### Microscopy

Light microscopy was performed with Axio Vert.A1 (Carl Zeiss) equipped with 100x oil immersion objective and AxioCam ICm1 camera (Carl Zeiss). Further, software Zeiss ZEN 2011 was used. Additionally, iScope IS.1153-EPLi (Euromex Microscopen) equipped with 100x oil immersion objective and Moticam X2 camera (Motic) was used including the software MotiConnect version 1.5.9.11.

### Genome Sequencing

Genomic DNA was isolated as described in (Hofmeyer *et al*. 2016). Shotgun libraries were generated using the Nextera XT DNA Sample preparation kit following the manufacturer instructions. The whole genome of *Y. lipolytica* strains 63, 1889 and H222*Δpox1-6* were sequenced with the Genome Analyzer IIx (Illumina, San Diego, USA) by the Göttingen Genomics Laboratory (G2L, http://appmibio.uni-goettingen.de/index.php?sec=g2l). The libraries were sequenced in a 112 bp paired end single indexed run. The sequencing datasets are available in the NCBI sequence read archive with the accession number PRJNA509765.

Paired reads were trimmed with BBDuk (version 37.25) using qtrim=rl, trimq=20, minlength=20. Then the reads were mapped against CLIB89 (Magnan *et al*. 2016) using the Geneious Mapper of Geneious 10.2.2 and the setting “Medium-Low Sensitivity/Fast” including iterative fine tuning (up to five times) to improve read alignment to INDEL regions. Additional settings for mapping were as follows: minimum mapping quality: 30, trimming of paired read overhangs was enabled and reads were only mapped if both reads of a pair mapped. The high mapping quality suppressed mapping against repetitive elements and therefore unused reads were mapped against single reference sequences. This compromised sequences of rDNA, mobile elements and a duplicated region on chromosome C of CLIB89 (Magnan *et al*. 2016).

For variant calling a minimum coverage with five reads and minimum variant frequency of 0.9 (with p-value ≦ 10^−6^) were set. Variant calling was not performed in regions with high coverage which were defined with coverage > (mean + 2*sdv).

Further analysis were done in R using tidyverse version 1.2.1. BLAST was performed in the fungi database of UniProtKB to find homologous sequences. But only reviewed records were selected. Homologous sequences were used to analyze the protein sequence variants by protein alignments using Geneious Alignment of Geneious 10.2.2 with BLOSUM62 matrix. Further, PROVEAN web server (v1.1.3) was used to predict the influence of the amino acid changes on protein functions using the default cutoff −2.5 (Choi *et al*. 2012).

## Results and Discussion

The productivity of *Y. lipolytica* was analyzed in numerous publications, revealing a substantial impact of process variables like pH and dissolved oxygen. These variables did not only have an impact on the yield but also on the product spectrum (Egermeier *et al*. 2017; Sabra *et al*. 2017). So far, the temperature was rarely considered and its effect on the formation of sugar alcohols has not been discussed in literature at all (Timoumi *et al*. 2018). The recommended growth temperature for *Y. lipolytica* is 25-30 °C and barely exceeds 34 °C (Barth and Gaillardin 1996). However, stable growth and product formation at higher temperatures would be beneficial especially for large scale processes as a reduction in cooling cuts energy costs. Further, thermal gradients that might appear in such processes can negatively impact fermentation performance if the strain is highly sensitive (Crater and Lievense 2018). Therefore we cultivated three *Y. lipolytica* strains, two wild type isolates (63 & 1889) and the modified lab strain H222*Δpox1-6* at 28 and 35 °C to learn about their behaviour in different media concerning growth, morphology, lipid accumulation and formation of citrate and sugar alcohols. Cultivations were performed in the complex medium YPD, the defined medium YNB and the minimal medium YSM. The first two media are commercially available and contain a balanced amount of carbon and nitrogen sources. They are broadly used for propagation of yeasts and with modified formulation for protein, lipid and organic acid production with *Y. lipolytica* (Madzak, Tréton and Blanchin-Roland 2000; Beopoulos *et al*. 2008; Gao *et al*. 2016). In this work both media contained 20 g L^−1^ glucose. The second minimal medium YSM triggers the lipid accumulation in *Y. lipolytica* due to its low nitrogen content as well as high glucose concentration (50 g L^−1^). Therefore it was used to analyze lipid production of the two wild type strains in comparison to H222Δ*pox1-6* harbouring a blockade of the fatty acid degradation (Gatter *et al*. 2014).

### Temperature- and medium-dependent growth

The cultivations were performed in shake flasks. At 28 °C the best growth regarding the maximal growth rate and OD_600_ values was achieved in the complex medium (YPD). The three strains differed notably in the achieved maximal OD_600_ (Fig. 1, Tab. S1 & S2). The strain 1889 reached the highest (OD_600_= 33) and the strain 63 the lowest OD (OD_600_= 18). The maximal growth rates and OD_600_ values dropped up to 70 % in the minimal media YNB and YSM. This was likely due to the low abundance or the complete absence of yeast extract and peptone, respectively, in the minimal media.

**Figure 1:**
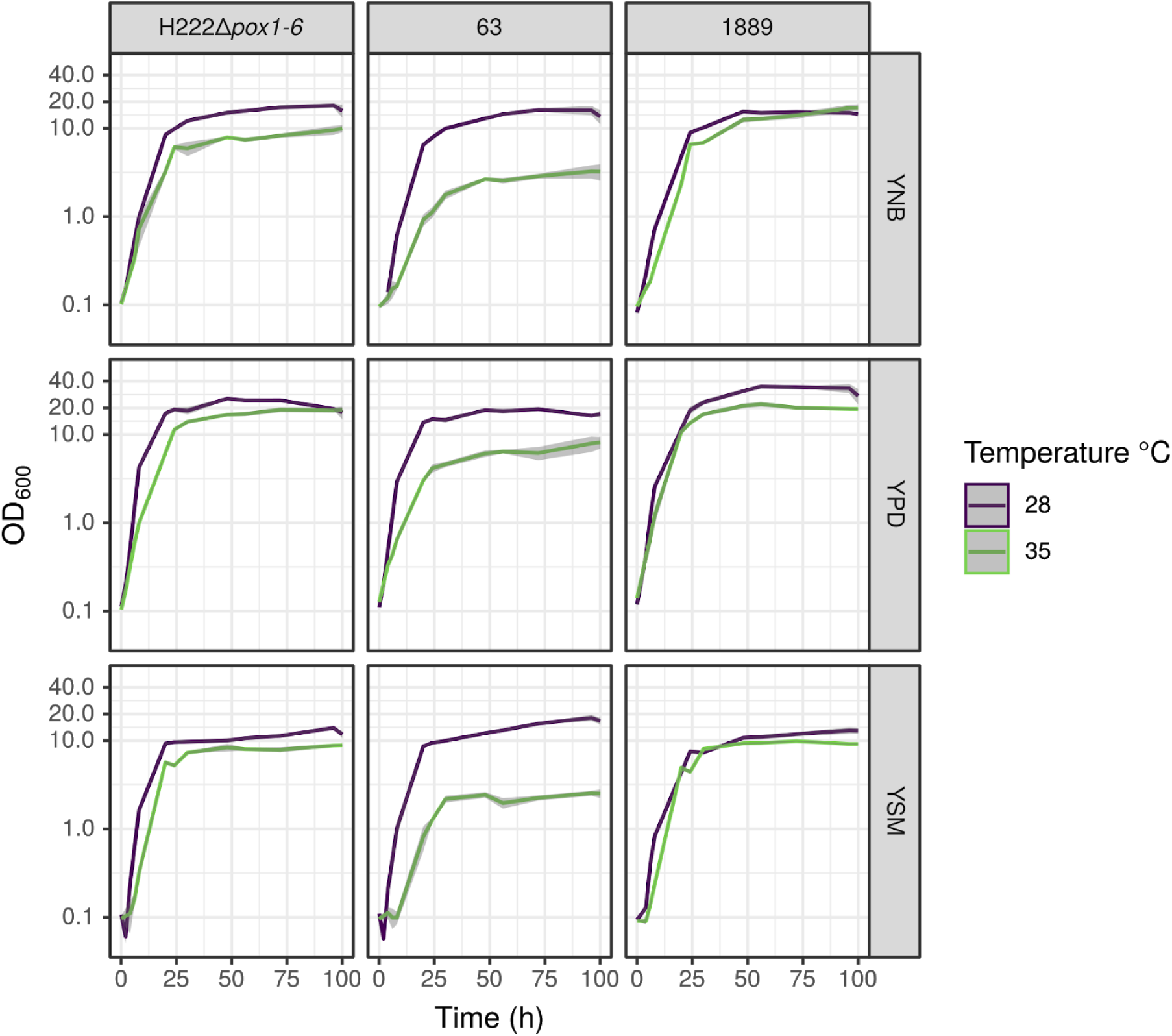
Growth profiles of the shake flask cultivation of *Y. lipolytica* strains H222Δ*pox1-6*, 63 and 1889 grown in the complex medium YPD and in the minimal media YNB and YSM, under different temperatures. Analysis was done with biological triplicates and the standard deviation is displayed as grey shadow.

In general, the temperature shift from 28 to 35 °C resulted in decreased growth for all three strains (Fig. 1). But the extent varied from 16 to 85 % reduction of the maximal growth rates and OD_600_ values despite two outliers where comparable or slightly higher values could be achieved (Tab. S1 & S2). This trend corresponds with the literature where the optimal growth temperature for *Y. lipolytica* is specified with 25 to 30 °C (Karasu-Yalcin, Bozdemir and Ozbas 2010; Papanikolaou, Chevalot, Komai *et al*. 2002). For H222, the ancestor strain of H222Δ*pox1-6*, the maximal growth rate exponentially increased from 24 to 30 °C and stagnated between 32 and 34 °C (Moeller *et al*. 2007). Nevertheless considerable growth could be measured for H222Δ*pox1-6* in YPD and for 1889 in YPD and YNB.

Similar to 28 °C the best growth was achieved in YPD followed by YNB and YSM. The strain 63 had serious growth deficits in all three media and reached only a maximal OD_600_ of 7.5 in YPD after 100 hours of cultivation. Taccari *et al*. determined for *Y. lipolytica* strain DiSVA C 12.1 that the negative impact of elevated cultivation temperatures on the biomass yield increases with higher substrate concentration (Taccari *et al*. 2012). This effect could not been seen in this work as the growth in YSM and YNB was similarly affected for 1889 and H222Δ*pox1-6* by the temperature shift.

### Temperature- and medium-dependent production of citrate and sugar alcohols

Optimal conditions for biomass formation not always match the prerequisite for optimal product yield. A classic example for this is the need for an high carbon to nitrogen ratio to trigger lipid accumulation in *Y. lipolytica* at the expense of the growth. The same applies for the production of citrate and sugar alcohols with *Y. lipolytica*. Next to the influence of the media composition we analyzed the influence of the elevated cultivation temperature on the formation of these metabolites in the *Y. lipolytica* strains 63, 1889 and H222Δ*pox1-6*. Therefore samples from the shake flask cultivations were analyzed with HPLC (Fig. 2).

**Figure 2:**
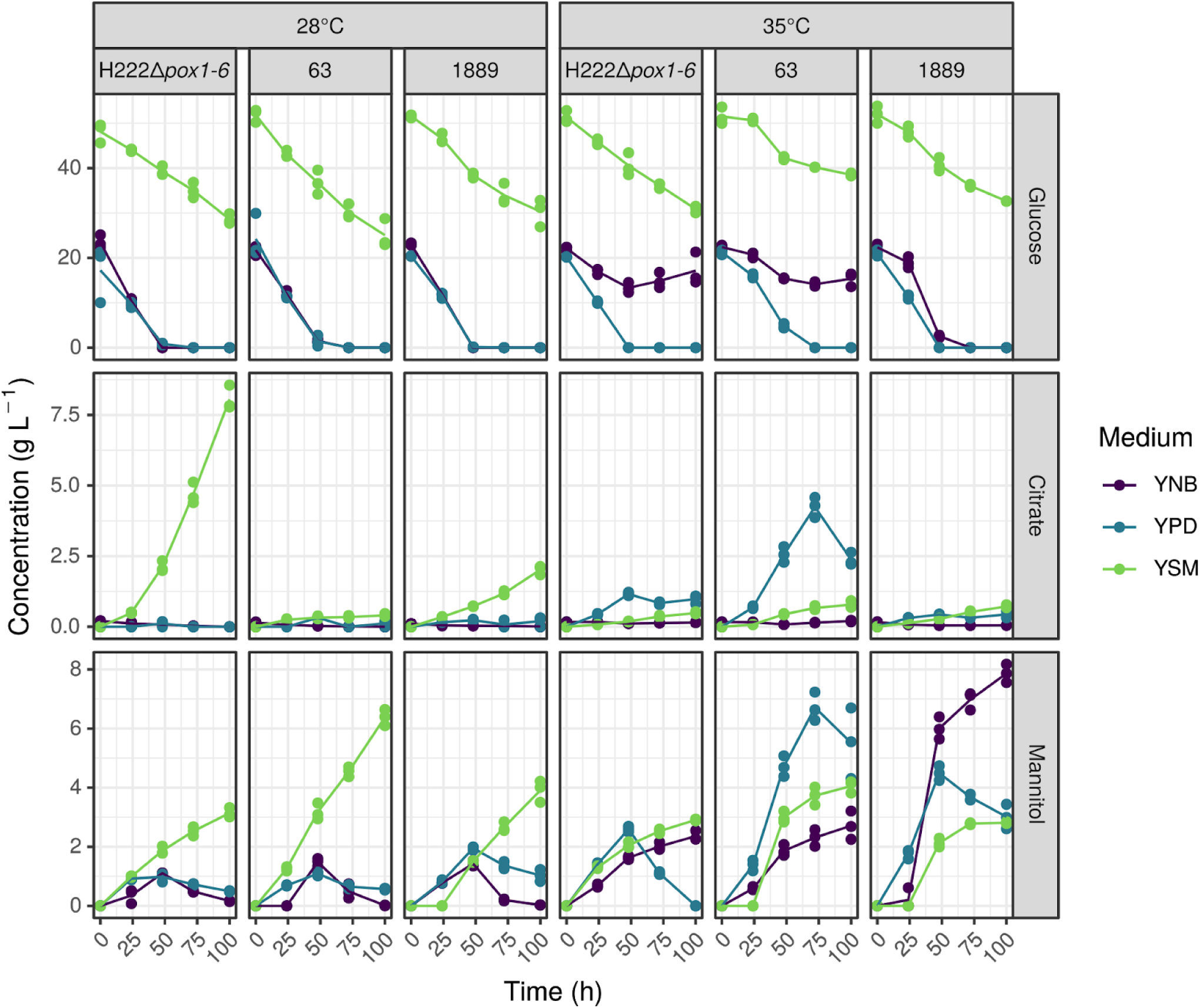
Consumption of glucose and formation of citrate and mannitol at 28°C and 35°C on different minimal and complex media for *Y. lipolytica* strains *H222Δpox1-6*, 63 and 1889. Biological replicates are displayed as single dots and graph lines represent the mean concentrations.

All three media contained glucose as carbon source, YPD & YNB 20 g L^−1^ and YSM 50 g L^−1^. The glucose consumption at 28 °C was similar for all three strains: In YPD and YNB the glucose was completely exhausted within 48 hours. In contrast, in YSM the strains were not able to convert the entire glucose. 30 to 25 g L^−1^ glucose remained after 100 hours of cultivation. Strain 1889 and H222Δ*pox1-6* which exhibited good growth performance at 35 °C, showed similar glucose consumption at 35 °C. Only in YNB H222Δ*pox1-6* stopped to take up glucose during the second half of the cultivation. This effect was also observed for *Y. lipolytica* 63. In addition, in YPD glucose exhaustion was delayed and in YSM 38 g L^−1^ glucose was left at the end of the cultivation of strain 63. The reduced glucose consumptions of H222Δ*pox1-6* and 63 matched the impaired growth at 35 °C. The high C/N ration of the YSM medium led to citrate and sugar alcohol formation in the three *Y. lipolytica* strains. H222Δ*pox1-6* accumulated citrate up to 8.1 and 1889 up to 2.0 g L^−1^. Considering the published data for *Y. lipolytica* the yields are quite low but it can be assumed that longer cultivation duration would had yielded higher values as the citrate concentration did not stagnate during the 100 hours (Abdel-Mawgoud *et al*. 2018). Furthermore, for the wild type of H222Δ*pox1-6* it was shown that extensive improvements of the cultivation conditions could significantly improve the yield up to 132 g L^−1^ citrate (Moeller et al. 2010). Only minor amounts of citrate could be detected for *Y. lipolytica* 63 at 28 °C. But surprisingly at 35 °C this strain produced up to 4.2 g L^−1^ citrate in the complex medium YPD, which was reused by the yeast cells after glucose exhaustion. This was the only case where significant citrate production could be observed at 35 °C. Neither H222Δ*pox1-6* nor 1889 accumulated more than 1 g L^−1^ citrate at this temperature.

The published results conclude that the optimal temperature for citrate production with *Y. lipolytica* usually is 28-30 °C and temperatures over 30 °C lead to reduced citrate concentrations (Morgunov, Kamzolova and Lunina 2013; Moeller *et al*. 2007; Karasu-Yalcin, Bozdemir and Ozbas 2010). For instance, the citrate concentration in a bioreactor fermentation of H222 dropped from 36.3 g L^−1^ at 28 °C to 13.5 g L^−1^ at 34 °C, a decline by 63 % (Moeller *et al*. 2007). In this work the shift to 35 °C in the shake flask cultivation of H222Δ*pox1-6* even led to 94 % reduction. Only 0.5 g L^−1^ citrate could be measured showing that elevated cultivation temperatures can almost completely suppress the citrate production. The induction of citrate formation by high temperatures as seen in this work for *Y. lipolytica* 63 has not described for this yeast in literature. Only for *Y. lipolytica* NBRC1658 it was shown that the maximal specific citrate production rate (g citrate (g cells)^−1^ h^−1^) raised with increasing temperature, yet the overall amount of citrate was decreased in comparison to lower temperatures (Karasu-Yalcin, Bozdemir and Ozbas 2010).

Next to citrate, the formation of the sugar alcohols arabitol, erythritol and mannitol was analyzed with HPLC. For both temperatures no arabitol accumulation could be detected and erythritol was produced only in low amounts, not exceeding 0.8 g L^−1^ [data not shown]. Mannitol formation could be measured for all three strains and media (Fig. 2). At 28 °C maximally 2.0 g L^−1^ have been produced in YPD and YNB as long as glucose was still present in the media. After glucose exhaustion mannitol was reused by the yeast cells. In contrast, in YSM mannitol was continuously produced reaching 6.4 g L^−1^ for *Y. lipolytica* 63 at 28 °C. With H222Δ*pox1-6* and 1889 only half as much was measured at this temperature. The mannitol production pattern was drastically altered at 35 °C. Most striking was the repeal of the requirement of high substrate concentrations to promote sugar alcohol formation. Mannitol production could be observed in the minimal medium YNB (containing only 20 g L^−1^ glucose and 5 g L^−1^ ammonium sulfate) and even in the complex medium YPD. But the medium preference was dependent on the strain, whereas for H222Δ*pox1-6* no clear medium preference could be defined. The highest values were reached by 63 with 6.7 g L^−1^ in YPD after 72 hours and by 1889 with 7.9 g L^−1^ in YNB after 100 hours. At the onset of glucose exhaustion the mannitol production was stopped and the sugar alcohol was re-consumed in most of the cases as already seen at 28 °C. Interestingly, only *Y. lipolytica* 1889 continued to accumulate mannitol in YNB beyond this point. This happened probably at the expense of cellular energy storages like glycogen or lipids (Sarris *et al*. 2011; Dulermo *et al*. 2015).

Also worthy of mention is the produced amount of mannitol despite the weak growth of *Y. lipolytica* 63 at 35 °C. It seems that the elevated cultivation temperatures shifted the carbon flux away from biomass production towards mannitol assimilation. The yield was significantly improved from 0.8 g (g CDW)^−1^ at 28 °C to 4.8 g (g CDW)^−1^ at 35 °C (Fig. S1).

The mannitol metabolism and its regulation in *Y. lipolytica* is only rudimentarily understood (Dulermo *et al*. 2015). However, it is assumed that sugar alcohols like mannitol and erythritol are accumulated by *Y. lipolytica* in response to high osmotic pressure, like high substrate or salt concentrations (Yang *et al*. 2014). Combination of high substrate concentrations with acidic pH further increases sugar alcohol contents (Egermeier *et al*. 2017; Yang *et al*. 2014). Moreover mannitol is also involved in the oxidative stress response reducing the level of reactive oxygen species especially hydroxyl radicals (Xu, Qiao and Stephanopoulos 2017; Sekova *et al*. 2018). Workman *et al*. also discussed a mannitol shuttle for *Y. lipolytica* which could serve as an alternative NADH recycling pathway under oxygen limitation (Workman, Holt and Thykaer 2013). The amount and product pattern depends on the strain and medium composition. In a comparative study higher polyole concentrations were reached with strains isolated from dairy products in comparison to common lab strains (Egermeier *et al*. 2017). Erythritol concentration can be influenced by addition of NaCl, pH change and the choice of carbon source, whereas mannitol accumulation seems to be only affected by high salt concentrations (Egermeier *et al*. 2017; Rymowicz, Rywińska and Marcinkiewicz 2009; Yang *et al*. 2014).

The factor temperature was so far not discussed for *Y. lipolytica* in the context of polyol biosynthesis. In contrast, glycerol accumulation in the model organism *Saccharomyces cerevisiae* has already been proven to promote resistance to heat, next to osmotic and oxidative stress (Chaturvedi, Bartiss and Wong 1997; Siderius *et al*. 2002). Obstruction of the glycerol biosynthesis in *S. cerevisiae* by deletion of the genes encoding for the glycerol-3-P dehydrogenase and glycerol-3-P phosphatase resulted in a temperature sensitive strain. Additionally, this defect could be abolished by addition of glycerol to the medium (Siderius *et al*. 2002). On the basis of our data it can be assumed that *Y. lipolytica* also specifically accumulates sugar alcohols in response to heat stress. At 28 °C all three strains produced only considerable amounts of mannitol when they were challenged by osmotic stress. In contrast, at 35 °C mannitol was also present in media with moderate substrate and salt concentrations. No additive effect could be observed if heat and osmotic stress were combined. Although the exact role of sugar alcohols in response of *Y. lipolytica* towards heat stress and its regulation needs to be further clarified, increased cultivation temperatures should be considered in future for optimization of the sugar alcohol production with *Y. lipolytica* especially on complex substrates like side streams.

### Temperature- and medium-dependent lipid accumulation

We analyzed the lipid amounts and fatty acid compositions of the shake flask cultivations after 100 hours incubation at 28 and 35 °C (Fig. 3). In contrast to the sugar alcohol production, the C/N ratio is also crucial for elevated cultivation temperatures as lipid contents over 20 % of the CDW could just be achieved in YSM, the only medium with a high C/N ratio. The highest lipid content was reached by H222Δ*pox1-6*. The lab strain with blocked fatty acid degradation accumulated 31.3 and 26.0 % of the CDW as lipids at 28 and 35 °C, respectively, demonstrating its robustness. The wild type isolates accumulated less amounts and showed contradictory trends by higher cultivation temperatures. The lipid content of strain 63 was halved in comparison to 28 °C whereas it was raised 1.3-fold for strain 1889. The high amount of lipids of strain 1889, hitherto genetically unmodified, along its slightly increased productivity at elevated temperatures highlight this strain as potential industrial host.

**Figure 3:**
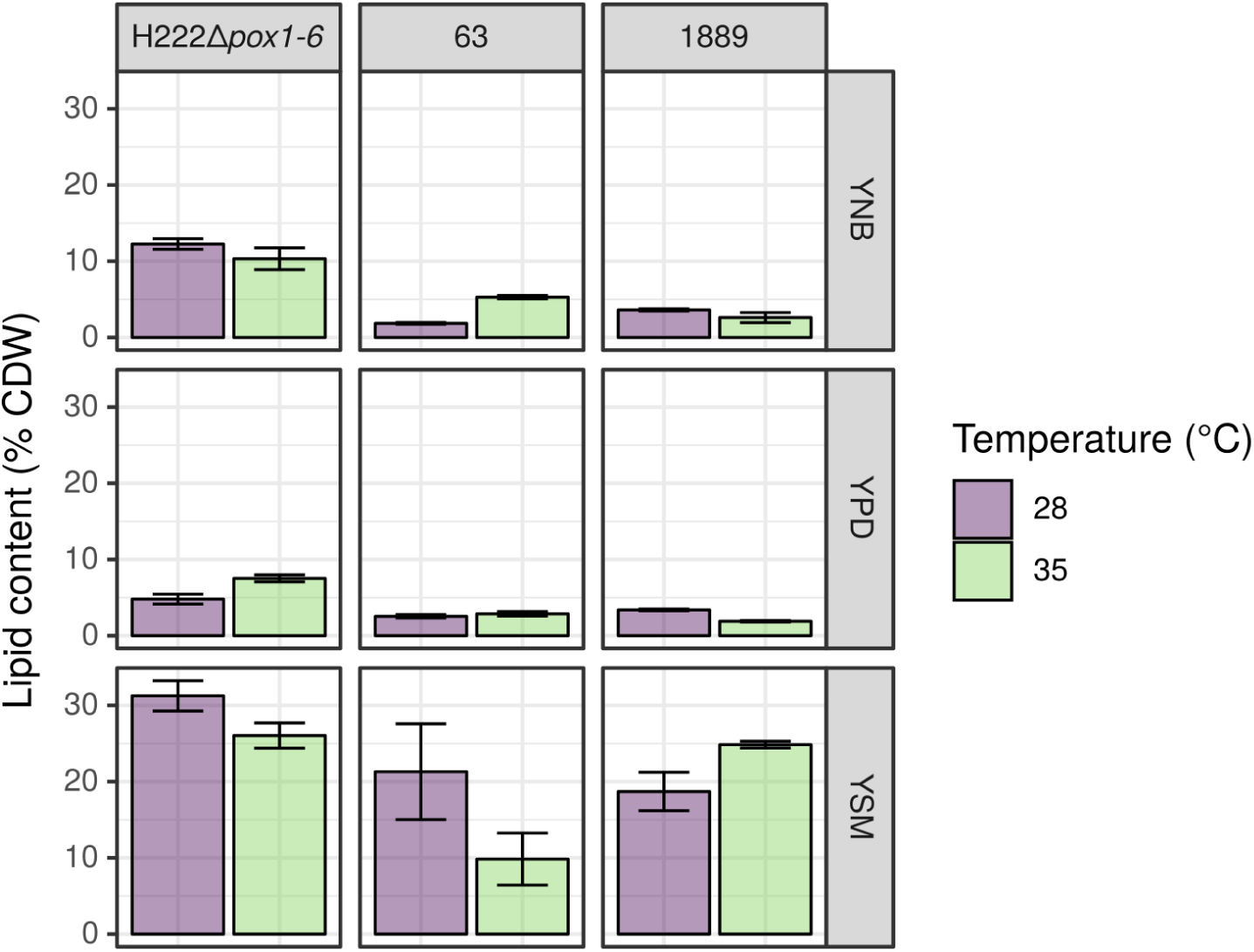
Lipid amounts of *Y. lipolytica* strains H222Δ*pox1-6*, 63 and 1889 after 100 hours of shake flask cultivation in the complex medium YPD and the minimal media YNB and YSM at two different temperatures.

Fatty acid compositions varied only slightly between the strains, media and temperatures and coincide with the published data for *Y. lipolytica* (Papanikolaou *et al*. 2009) (Fig. S2A). The most dominant fatty acids in all samples were oleic and stearic acid. No clear trend regarding the ratios of unsaturated to saturated fatty acids could be observed between the cultivation temperatures (Fig. S2B). In contrast, at lower temperatures (10-12 °C) the degree of unsaturation slightly increases for *Y. lipolytica* (Ferrante, Ohno and Kates 1983; Tezaki *et al*. 2017). Beyond that, the influence of the temperature on the lipogenesis was rarely discussed in literature for this yeast (Carsanba, Papanikolaou and Erten 2018). A comparable trend as seen in this work for the lab strain H222Δ*pox1-6* and the wild type isolate 63 is also described for *Y. lipolytica* ACA-CD 50109 grown on stearin. The optimal temperature for lipid production with this strain was 28 °C and incubation at 33 °C led to reduced lipid yields (Papanikolaou I. Chevalot M. Komai *et al*. 2002). An increase of the lipid content due to higher cultivation temperatures is to our knowledge so far not published.

### Temperature- and medium-dependent cell morphology

Dimorphic transition of *Y. lipolytica* is thought to be part of the adaptation to environmental fluctuations and can be induced by a variety of external factors like pH, temperature, carbon and nitrogen source, reviewed in (Timoumi *et al*. 2018). Within the reported data contradictions arose, suggesting a strain specific response (Timoumi *et al*. 2018). Therefore we analyzed the cellular morphology of *Y. lipolytica* H222Δ*pox1-6*, 63 and 1889 during the shake flask cultivations. Samples were analyzed by microscopy and flow cytometry. The later is based on the forward scatter (FSC) detection and provides an approximation of the cell sizes (Shapiro 2005). Whereas increasing signals in the FSC hint formation of pseudohyphae or increased cell size.

At 28 °C yeast like cell shape was predominant (Fig. 4B). But the strains trend to decreased cell size in YSM and YPD in the course of the cultivation (Fig. 4A). An opposite effect could be seen for YNB. Both trends are not only independent of the strain but also of the different extent and timepoints of the C source exhaustion. Makri *et al*. reported a morphological change based on the metabolic activities during different growth phases. Large cells were predominant in the lipogenic phase and small cells during citrate production (Makri, Fakas and Aggelis 2010). But the cause for changed cell sizes in this work remains unclear.

**Figure 4:**
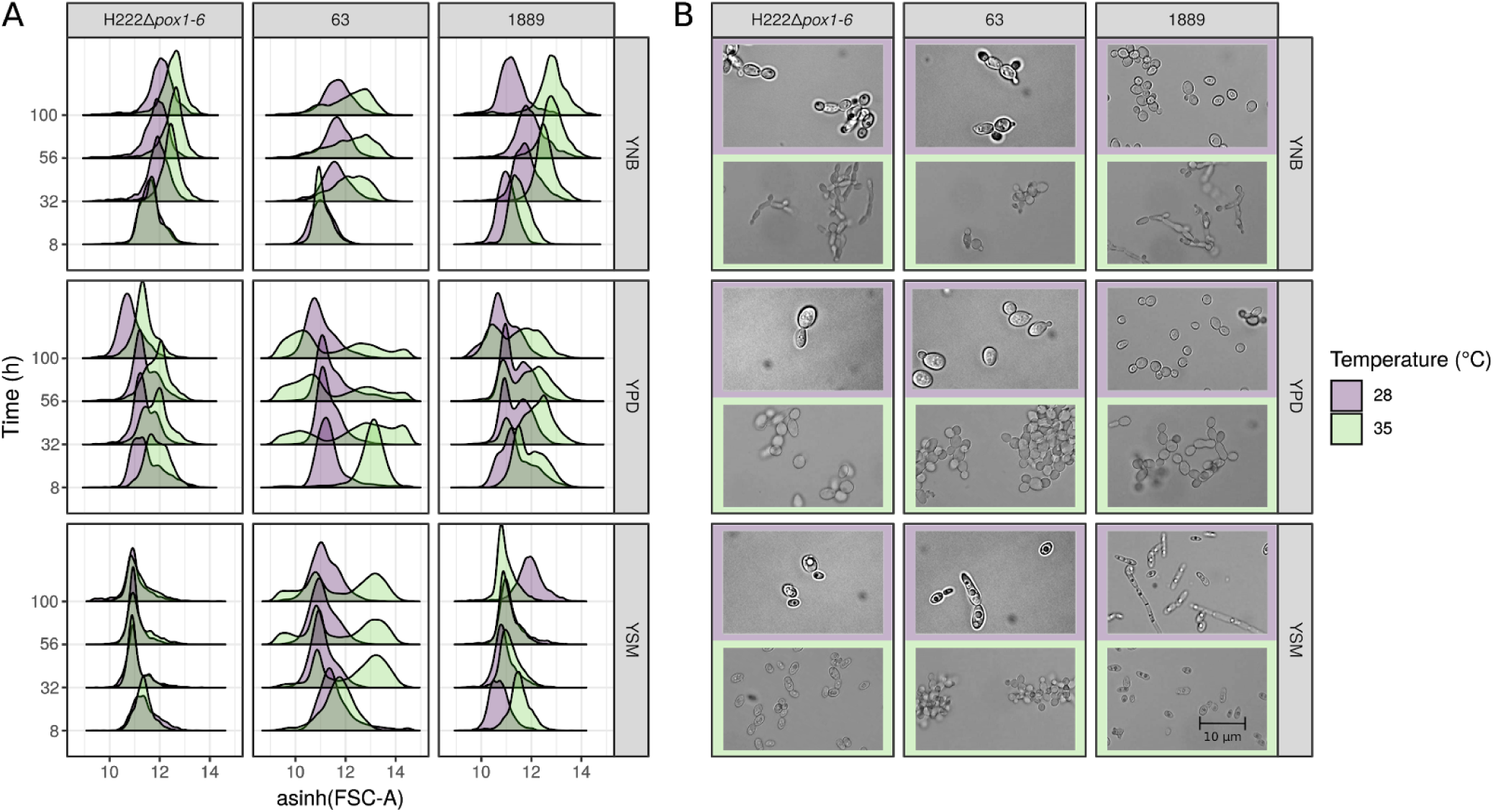
Flow cytometry data of *Y. lipolytica* strains H222Δ*pox1-6*, 63 and 1889 at two temperatures and in three different media. **A:** Shown are the inverse hyperbolic sine transformations of forward scatter area. **B:** Pictures show microscopy at t = 56 h.

Heat stress appears to evoke different physiological reactions depending on the strain, which is in line with the varying temperature tolerance of the strains observed in the preceding analysis. *Y. lipolytica* 63 had the strongest reaction to the elevated temperature and formed cell agglomerates (Fig. 4B). The agglomerates differed in size in dependency to the medium and consisted of small cells with a yeast like cell shape. This morphology is reflected in the diffuse spreading of the FSC signals as seen in figure 4A. The cell clumps were also observed for H222Δ*pox1-6* and 1889 in YPD. In contrast, in YNB both strains appeared as small branched pseudohyphae (Fig. 4B) resulting in slightly increased FSC signals (Fig. 4A). Formation of pseudohyphae due to heat stress is in accordance with a previous publication (Kawasse *et al*. 2003) whereas the detection of cell agglomerates seem to be absent in literature for *Y. lipolytica*.

The fewest phenotypical changes were observed for H222Δ*pox1-6* and 1889 in YSM, very likely due to the pre-cultivation in the same medium at 28 °C. As seen in figure 2 the yeast cells accumulate mannitol in YSM at this temperature. It can therefore be assumed that the cells were already adapted to a stressful environment when they were used as an inoculum for the main cultures at 35 °C. Interestingly this adaptation was not sufficient for *Y. lipolytica* 63. The phenomenon of cross-resistance observed here is already described for *Y. lipolytica* as well as for *S. cerevisiae* and other eu- and prokaryotic organisms (Święciło 2016). Mild pretreatment with a stressor led to increased viability of the yeast cells when they were exposed to another stressor. For instance, incubation of *Y. lipolytica* with 0.5 mM H_2_O_2_ enhanced the survival rate at 45 °C by 30 % in comparison to untreated cells (Biryukova *et al*. 2007).

The presented flow cytometry method seems to be suitable to make assumptions about the fitness of different *Y. lipolytica* strains in respect to growth in a rapid manner. Strain H222Δ*pox1-6* which reacted robust under both temperature regimes remained stable in its scattering peak area over time. The strong reaction of strain 63 is as well reflected in the plot of the cytometry data. A principal component analysis (Fig. S3) indicates that the detected change in cell size is responsible for these variations. No assumptions about secreted metabolites or lipid production can be made from the cytometric data.

### Analysis of Genotypes

Shotgun sequencing was performed to gain deeper insight into the genetic background of the three different phenotypes. The resulting raw reads were mapped against the annotated genome of CLIB89 as described in Materials & Methods. Good sequencing depths were achieved for H222Δ*pox1-6* and 1889 with 66x and 61x, respectively, and 98 % of the reference genome was covered with at least five reads (threshold for variant calling) for both strains (Tab. S3). Despite a second sequencing attempt, only a relatively low amount of reads could be sequenced for strain 63. In consequence sequencing depth was substantially lower (22x) in comparison to 1889 and H222Δ*pox1-6* and only 85 % of the reference genome was covered with enough reads to perform variant calling. Reads were also used to predict the occurence of mobile elements in the wild type strains (Tab. S4). Non-LTR LINE retrotransposon Ylli which exists in several copies in CLIB89 and CLIB122 (Magnan *et al*. 2016) was also found in all three strains. Strain 1889 additionally contained the Ty3/Gypsy LTR retrotransposon Tyl3. Further retroelements which were identified in other *Y. lipolytica* strains, like Ylt1 and Tyl6 (Schmid-Berger, Schmid and Barth 1994; Kovalchuk *et al*. 2005), were absent in our strains. Mutator-like DNA transposon Mutyl was only partially present in the three strains. But as already described for H222, the genome of H222Δ*pox1-6* contained the DNA transposon Fotyl (Gaillardin, Mekouar and Neuvéglise 2013).

In *Y. lipolytica* 1889 no reads could be found for a 10.5 kb region on chromosome B containing YALI1_B10231g, YALI1_B10263g and YALI1_B10288g. The first gene encodes a Acyl/Aryl-CoA-ligases (Aal) which is involved in fatty acid activation in peroxisomes among nine other Aal proteins (Dulermo *et al*. 2016). The protein encoded by YALI1_B10288g is similar to Opt2 of *S. cerevisiae*. *Sc*Opt2 is necessary for the glutathione redox homeostasis and plays an important role in the adaptation to altered lipid asymmetry in the plasma membrane in this yeast (Elbaz-Alon *et al*. 2014; Yamauchi *et al*. 2015). Its function in *Y. lipolytica* was not characterized so far. No annotation is given for YALI1_B10263g. But according to InterPro it contains a reverse transcriptase and Rnase H-like domain suggesting that it encodes a mobile element or a remnant of it.

The three strains exhibited three nucleotide polymorphisms/kb on average and about 10 % of them caused a sequence variation on protein level resulting in up to 6608 protein polymorphisms per strain (Fig. 5A). The vast majority were amino acid substitutions and in total 9926 different protein polymorphisms were found in the three strains (Fig. 5B). Comparison between them suggest that strain 63 and H222Δ*pox1-6* were more closely related to each other as to 1889 since they shared 44 % of all identified protein polymorphisms (Fig. 6). In contrast, strain 1889 had the highest proportion of unique polymorphisms.

**Figure 5:**
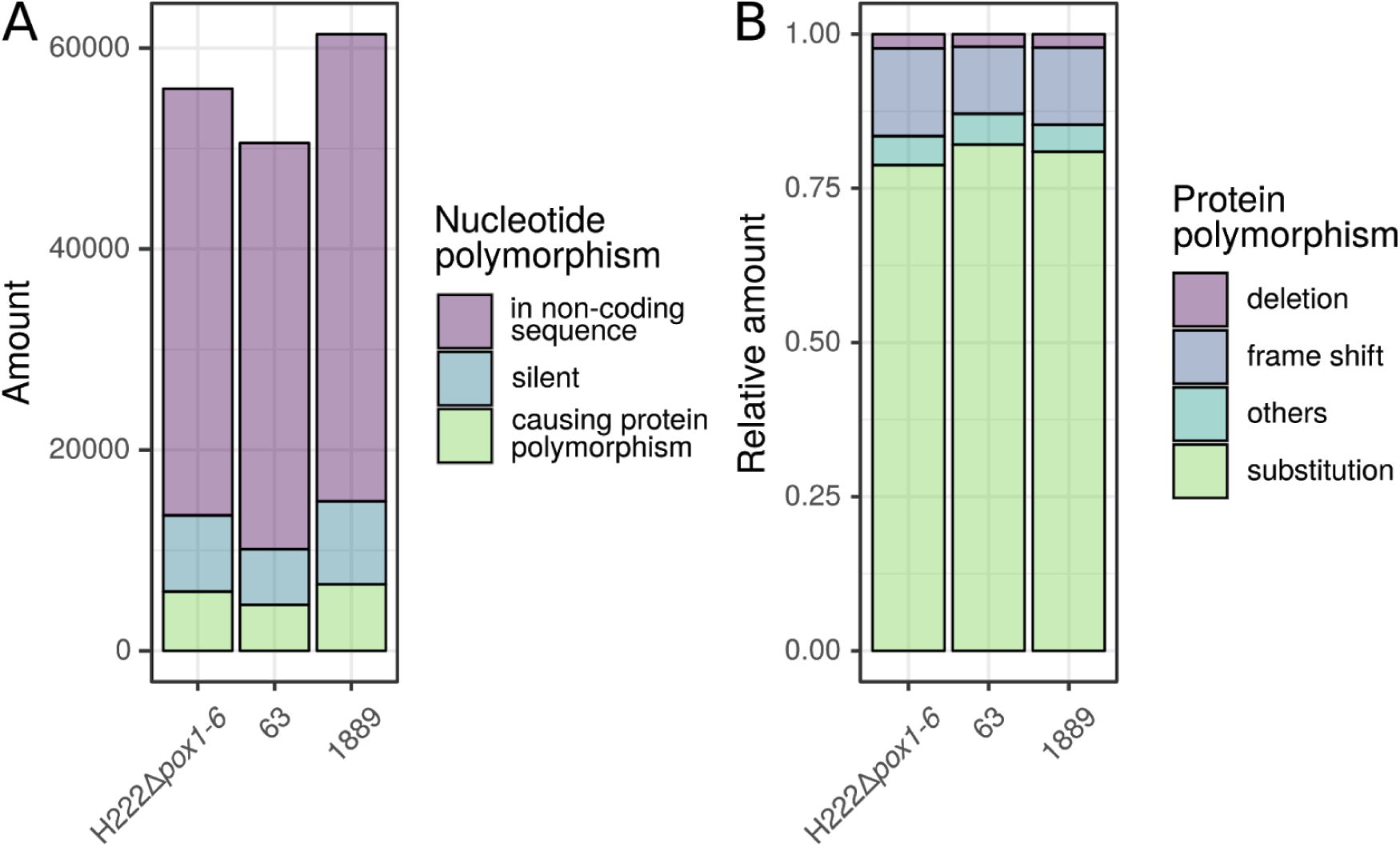
Count and type of polymorphism found in *Y. lipolytica* strains H222Δ*pox1-6*, 63 and 1889 in comparison to the reference strain CLIB89. **A:** Counts of predicted silent, mis- or nonsense mutations. **B:** Type and proportion of protein polymorphism. Counts for insertion, extension, start codon loss and truncation are summed up as “others”.

**Figure 6:**
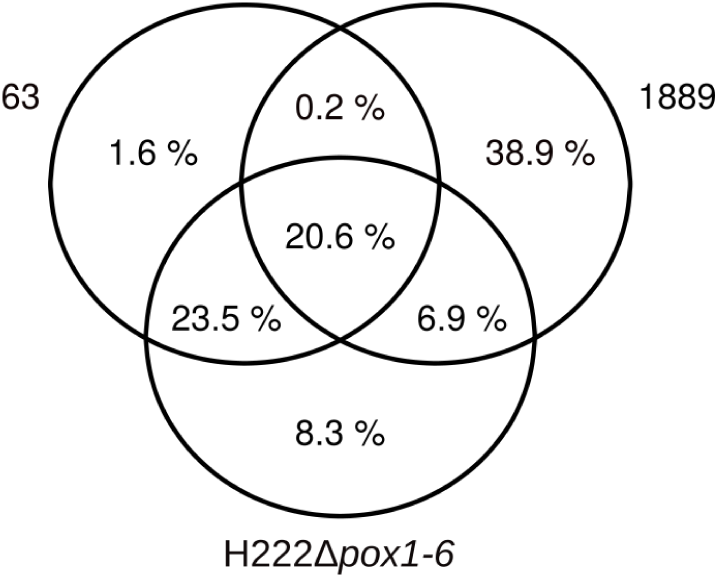
Inter-strain comparison of the protein polymorphisms. In total 9926 different protein polymorphisms were found and the Venn diagram shows the proportion of shared protein polymorphisms.

The library of identified protein polymorphisms was screened for potential candidates that may be responsible for the different phenotypes. Therefore, identical sequence variants which appeared in all three strains or that occured at loci which were not covered in all three strains with at least 5 reads were rejected. The filtered dataset contained 2757 affected genes compromising 34 % of all annotated genes of CLIB89. In addition to the annotations of the CLIB89 genome, the entries in the KEGG and Gene Ontology databases for *Y. lipolytica* were used to screen the data. But unfortunately, no function or homology was assigned to 42 % of the affected genes.

The dataset was screened for genes known to be involved in lipid metabolisms (Beopoulos *et al*. 2009; Dulermo *et al*. 2016; Silverman *et al*. 2016; Seip *et al*. 2013; Kerkhoven *et al*. 2016), dimorphism (Gonzalez-Lopez *et al*. 2002; Pomraning *et al*. 2018), mannitol formation (Dulermo *et al*. 2015) and stress response (Biryukova *et al*. 2007; Biryukova *et al*. 2008; Flores, Gancedo and Petit 2011; Yang *et al*. 2015) in *Y. lipolytica*. Additionally, proteins of MAPK cascades were checked. MAPK cascades are signaling pathways commonly found in eukaryotic cells which are involved, inter alia, in cell growth and adaptation to stress and they regulate the expression of various genes in response to extracellular signals (Chen and Thorner 2007; Morano, Grant and Moye-Rowley 2012; Dunayevich *et al*. 2018). However, in comparison to *S. cerevisiae*, MAPK cascades are only studied to some extent in *Y. lipolytica* (Cervantes-Chávez *et al*. 2009; Rzechonek *et al*. 2018).

Sequence variants for several proteins were identified (Tab. S5 & S6), whereby single amino acid substitutions were predominant as already seen before. Only one case of a premature stop codon was found in YALI1_C29447g in strain 1889. The encoded protein is not chararized but it shares 45 % similarity with the with osmosensing histidine protein kinase Sln1 of *S. cerevisiae*. However, we could not observe a clear phenotypic difference between 1889 and H222Δ*pox1-6* regarding the response to high osmolarity (growth, fitness and polyole formation). Either the function of YALI1_C29447p differs from *Sc*Sln1 or its lack is compensated by YALI1_F12254g which also encodes a protein with 46 % similarity with *Sc*Sln1. Alternatively the osmolarity could also be sensed by other proteins as seen for *S. cerevisiae* (O’Rourke and Herskowitz 2002).

As most of the affected proteins are not characterized in *Y. lipolytica* sequence, alignments with homologous from *S. cerevisiae* and other yeasts were performed to investigate the effects of the amino acid substitutions and indels on protein functions (see material and methods section). Unfortunately, due to missing assignments of affected residues or incomplete characterization of homologous genes, manual alignments did not lead to compelling results. Therefore a widely-used prediction tool, the PROVEAN web server, was used for further analysis of the variants (Choi *et al*. 2012). On the basis of pairwise alignments with homologous and distantly related sequences, this software predicts the damaging effect of variations (Tab. S6). For strain 63 deleterious amino acids substitutions were found for the MAP kinase *Yl*Ste11 (YALI1_F18202p) and YALI1_B20634p. The later is homologous to the histidine kinase Chk1 of *C. albicans* (49 % similarity). Deletion of Ste11 in *Y. lipolytica* abolishes mycelial growth (Cervantes-Chávez and Ruiz-Herrera 2006) and Chk1 null mutants of *C. albicans* flocculate under certain cultivation conditions (Calera and Calderone 1999). Both phenotypes coincide with the observed formation of cell agglomerates consisting of yeast-like cell for strain 63. On contrary, the data do not give any hints for genotypic basis of the temperature sensitivity of this strain. Interestingly, several deleterious sequence variations were found for 1889 and but none for H222Δ*pox1-6*, despite their similar phenotypes. However, genotypic diversity does not mandatory lead to distinct phenotypes (Wagih *et al*. 2018) as well as the prediction accuracy of such tools is limited and dependent on the amount of available homologous sequences (Choi *et al*. 2012).

In both cases, manual or automated prediction, the identified sequence variants must be verified in further studies *in vivo* to determine their relevance.

### Summary

In this work, we characterize three *Yarrowia lipolytica* strains in respect to their growth rate, key metabolites and lipid accumulation capabilities during normal (28 °C) and high temperatures (35 °C) in different media. Additionally the morphology during these cultivation conditions is determined using flow cytometry and microscopy. We could observe increased production of sugar alcohols at elevated temperatures which was not described so far for *Y. lipolytica*. Especially wild type strain 1889 showed a robust growth and lipid formation at elevated temperatures. The described flow cytometry method allowed for the rapid detection of varying cell size resulting from cell stress. In contrast to the common formation of pseudohyphae in stressful environments we observed formation of agglomerates with yeast-like cell for the wild type strain 63. In order to correlate the observed phenotypes with possible genotypes draft genome sequencing was performed but only partially yielded conclusive results. Correlation of the observed phenotypes with sequenced genomes was impeded by the large number of affected proteins and the low quality of available information about sequence-function relationships of these proteins. Nevertheless the obtained genome data are a valuable basis for further characterization and modification of these strains.

## Funding

This work was supported by the Fachagentur für Nachwachsende Rohstoffe [FKZ: 22007413 to SH, FB, SB] and by the by the Hessian Ministry for Science and Art through the LOEWE-project CompuGene to JK. The authors have declared no conflicts of interest.

## Authors’ Contribution

SH, SB & FB performed the cultivations whereby flow cytometry analysis was done by FB, HPLC analysis by SB and lipid analysis by SH. Shotgun sequencing was conducted by RD and AT, SH analyzed the relating data with aid of FB. SH drafted the manuscript. All authors discussed the results and contributed to the final manuscript.

## Acknowledgements

JK would like to thank Frieder Schauer and Anne Reinhard for their continued support.

## Supplementary Materials

**Table S1:** Maximal growth rates µ_max_ of the shake flask cultivations of *Y. lipolytica* strains H222Δ*pox1-6*, 63 and 1889.

| | | $\mu_{\max}$ (h<sup>-1</sup>) | | | | | | | | |
| --- | --- | --- | --- | --- | --- | --- | --- | --- | --- | --- |
| Temp. | Medium | H222Δpox1-6 | | | 63 | | | 1889 | | |
| 28 °C | YNB | 0.58 | ± | 0.02 | 0.48 | ± | 0.05 | 0.57 | ± | 0.00 |
|  | YPD | 1.27 | ± | 0.15 | 0.93 | ± | 0.04 | 1.20 | ± | 0.08 |
|  | YSM | 0.66 | ± | 0.07 | 0.40 | ± | 0.03 | 0.35 | ± | 0.02 |
| 35 °C | YNB | 0.27 | ± | 0.02 | 0.07 | ± | 0.01 | 0.34 | ± | 0.02 |
|  | YPD | 0.68 | ± | 0.03 | 0.18 | ± | 0.01 | 0.94 | ± | 0.06 |
|  | YSM | 0.41 | ± | 0.02 | 0.18 | ± | 0.01 | 0.41 | ± | 0.02 |

**Table S2:** Maximal optical densities OD_600 max_ reached in the shake flask cultivations of *Y. lipolytica* strains H222Δ*pox1-6*, 63 and 1889.

|  |  | OD<sub>600 max</sub> |  |  |  |  |  |  |  |  |
| --- | --- | --- | --- | --- | --- | --- | --- | --- | --- | --- |
| Temp. | Medium | H222Δpox1-6 | | | 63 | | | 1889 | | |
| 28 °C | YNB | 16.7 | ± | 1.3 | 15.1 | ± | 1.6 | 15.1 | ± | 0.1 |
|  | YPD | 22.0 | ± | 0.7 | 17.8 | ± | 0.7 | 32.6 | ± | 2.4 |
|  | YSM | 11.4 | ± | 0.6 | 16.2 | ± | 1.4 | 12.4 | ± | 1.0 |
| 35 °C | YNB | 8.9 | ± | 0.6 | 3.2 | ± | 0.6 | 16.8 | ± | 1.6 |
|  | YPD | 18.5 | ± | 1.0 | 7.5 | ± | 1.3 | 20.5 | ± | 0.8 |
|  | YSM | 8.4 | ± | 0.4 | 2.4 | ± | 0.1 | 9.5 | ± | 0.2 |

**Figure S1:**
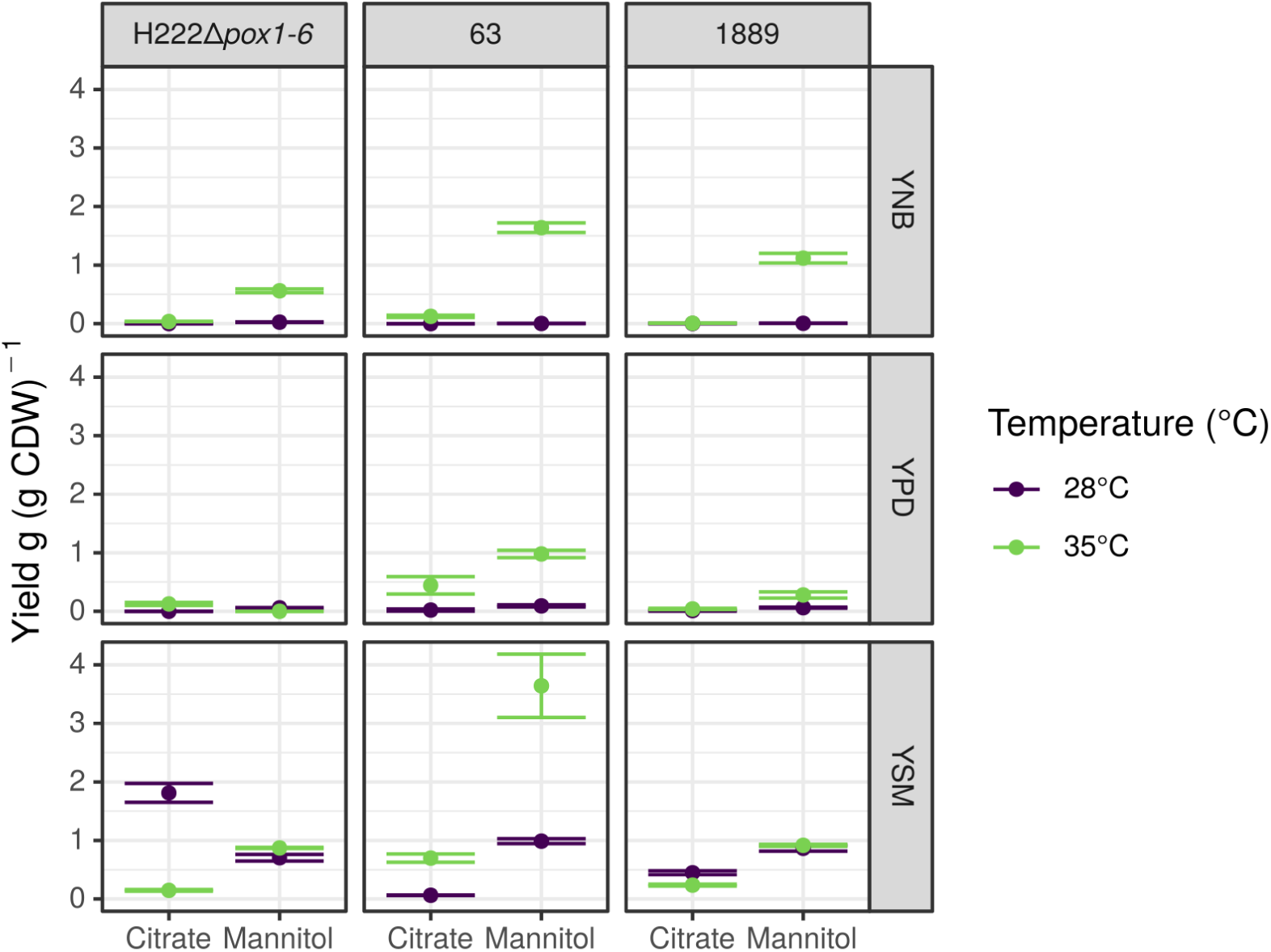
Yields of the citrate and mannitol formation with *Y. lipolytica* strains H222Δ*pox1-6*, 63 and 1889 cultivated in different media. Cultivation at elevated temperature significantly increased the mannitol yield of *Y. lipolytica* 63 in YSM.

**Figure S2:**
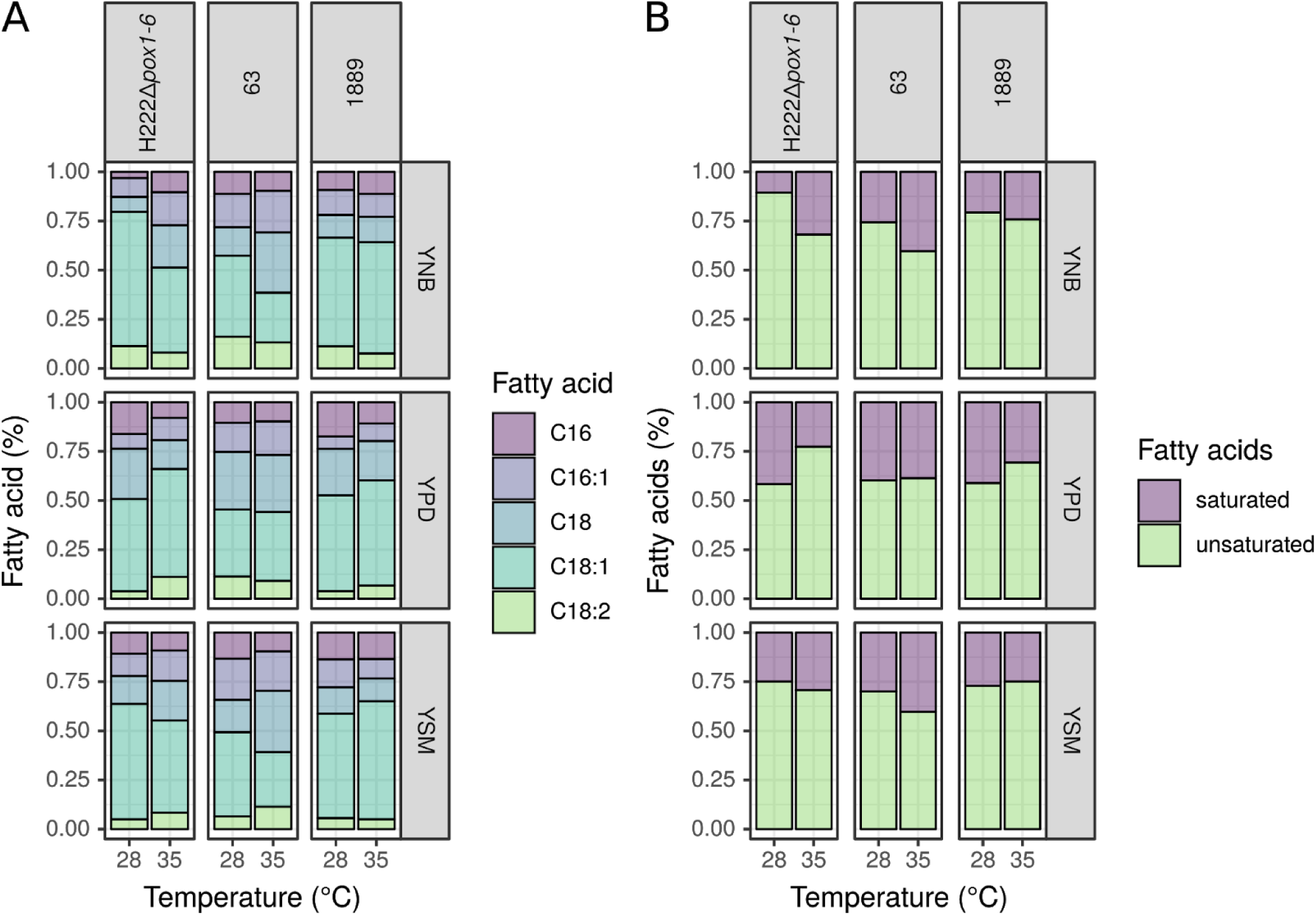
Fatty acid compositions of *Y. lipolytica* strains H222Δ*pox1-6*, 63 and 1889 after 100 hours of shake flask cultivation in the complex medium YPD and the minimal media YNB and YSM at two different temperatures.

**Figure S3:**
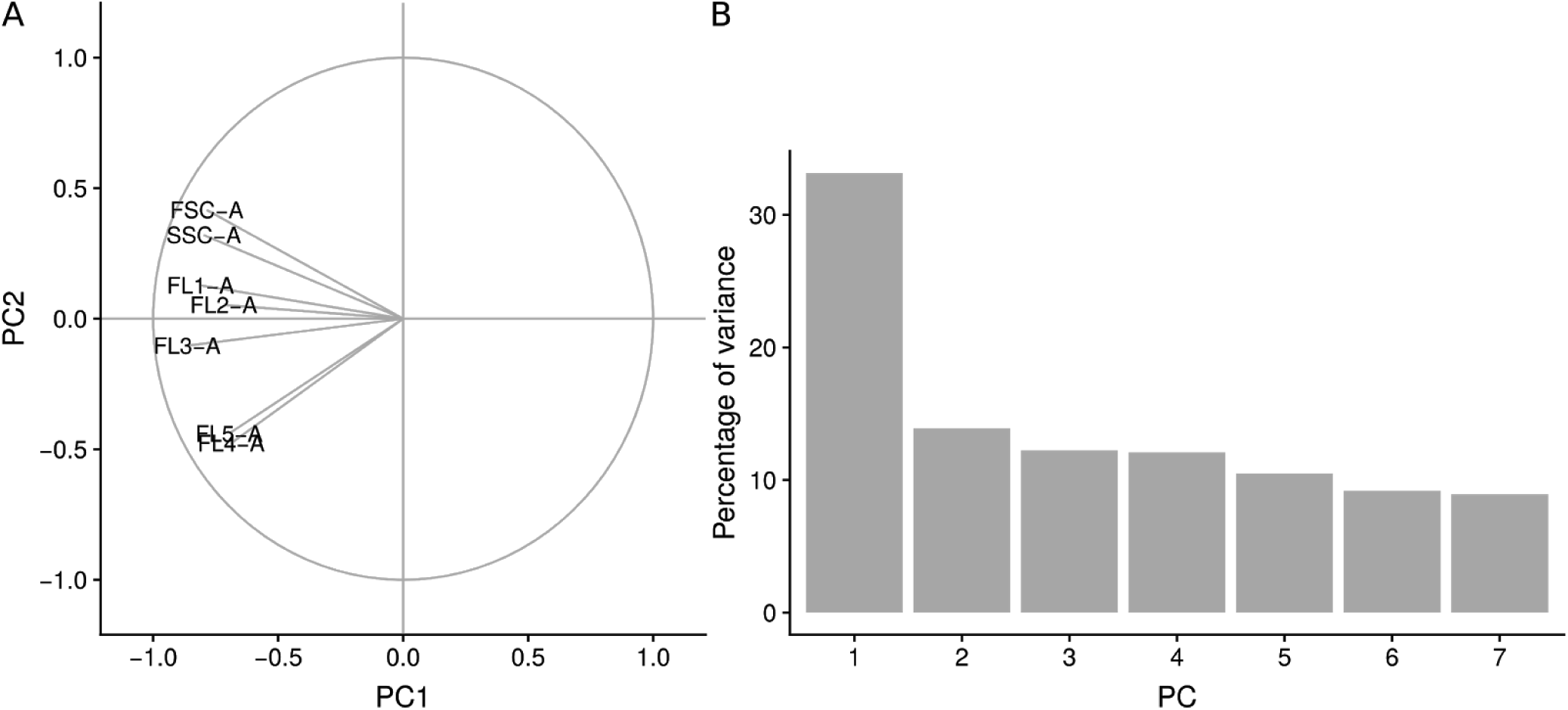
Principal component analysis of all fluorescence and scattering channels measured by flow cytometry during the course of *Y. lipolytica* cultivation using strains H222*Δpox1-6*, 63 and 1889 in the media YPD, YSM and YNB as shown in figure 4. Multicollinearity between channels is strong, indicating that independently of all measured variables, cell size appears to be mainly responsible for variation measured in all channels. Fluorescence was measured with the following channels, filters and corresponding PMT settings: FL1-A (525/50): 50.0 %, FL2-A (585/30): 40.0 %, FL3-A (617/30): 58.5 %, FL4-A (665/30): 40.0 %, FL5-A (720/60): 40.0 %, FL6-A (785/60): 40.0 %, FSC-A: 1, SSC-A: 17.0 %

**Table S3:** Shotgun sequencing of *Y. lipolytica* H222Δ*pox1-6*, 63 and 1889. Reads were mapped against the annotated genome of strain CLIB89 and variant calling was performed as described in Materials and Methods.

| | H222Δpox1-6 | 63 | 1889 |
| --- | --- | --- | --- |
| **Reads** |  |  |  |
| total reads | 17391232 | 6260112 | 16546422 |
| mapped reads | 92 % | 89 % | 91 % |
| mapped reads (incl. repetitive elements) | 96 % | 96 % | 96 % |
| **Depth of coverage** |  |  |  |
| mean | 66 | 22 | 61 |
| sdv | 63 | 37 | 52 |
| low coverage (< five reads) | 2.2 % | 14.9 % | 2.3 % |
| high coverage (> mean+2*sdv) | 0.03 % | 0.03 % | 0.04 % |
| **Variant calling** |  |  |  |
| identified nucleotide polymorphisms | 55955 | 50550 | 61378 |
| identified protein polymorphisms | 5886 | 4559 | 6608 |
| CDS with protein polymorphisms | 2873 | 2428 | 3161 |

**Table S4:** Mobile elements in *Y. lipolytica* H222Δ*pox1-6*, 63 and 1889. Reads were mapped against reference sequences (GenBank accessions nummers are given in brackets) to predict their occurrence. Plus sign means that at least 90 % of the reference sequence was covered.

|  | **H222Δ*pox1-6*** | **63** | **1889** |
| --- | --- | --- | --- |
| **Retrotransposons** |  |  |  |
| Ylt1 (AJ310725) | - | - | - |
| Tyl3 (CP017555) | - | - | + |
| Tyl6 (AJ746250) | - | - | - |
| Ylli (AJ319752) | + | + | + |
| **DNA transposons** |  |  |  |
| MutA* (AJ621548) | + | + | + |
| Fotyl (AJ745091) | + | - | - |

\* first 269-537 nt of CDS were not covered

**Table S5:** Genes with protein polymorphisms for *Y. lipolytica* strains H222Δ*pox1-6*, 63 and 1889 encoding for proteins involved in heat shock response, dimorphism and signal transduction of stress. Sequence variants which were identical in all three strains were not considered.

| Gene | Annotation (UniProtKB entry) | Function |
| --- | --- | --- |
| **Response to heat shock** |  |  |
| YALI1_C10555g | similar to *C. albicans* CaHSC82 (P46598) | heat shock protein |
| YALI1_D26051g | similar to *S. cerevisiae* HSP12 (P22943) | heat shock protein |
| YALI1_E19871g | similar to *S. pombe* DNAJ (Q9UTQ5) | HSP binding |
| YALI1_E22218g | similar to *S. cerevisiae* HSP42 (Q12329) | heat shock protein |
| YALI1_E36568g | similar to *S. cerevisiae* TSL1 (P38427) and TPS3 (P38426) | trehalose metabolism |
| **Signal transduction of stress (MAPK cascade)** |  |  |
| YALI1_B20634g | similar to *C. albicans* CHK1 (O59892) | MAPK cascades |
| YALI1_C29447g | similar to *S. cerevisiae* SLN1 (P39928) | MAPK cascades |
| YALI1_D00663g | similar to *S. cerevisiae* MSB2 (P32334) | osmosensor |
| YALI1_E03789g | similar to *C. albicans* CaNIK1 (O42696) | MAPK cascades |
| YALI1_E40089g | similar to *S. cerevisiae* SLT2 (Q00772) | MAPK cascades |
| YALI1_F12365g | similar to *S. cerevisiae* BCK1 (Q01389) | MAPK cascades |
| YALI1_F17657g | similar to *S. cerevisiae* LRG1 (P35688) | signal transduction |
| YALI1_F18202g | YISTE11 (Q7Z8J5) | MAPK cascades |
| YALI1_F19635g | similar *S. cerevisiae* RIM15 (P43565) | protein kinase |
| YALI1_F22241g | similar to *S. cerevisiae* SPA2 (P23201) | MAPK cascades |
| **Sugar alcohols** |  |  |
| YALI1_E15452g | similar to *Candida* sp. HA167 XDH (O74230) | putative mannitol dehydrogenases |

| **Lipid metabolism** |  |  |
| --- | --- | --- |
| YALI1_B10231g* | similar to *N. crassa* related to 4-coumarate-CoA ligase (Q7SDW1) | fatty acid activation |
| YALI1_B22375g | similar to *S. cerevisiae* ERG2 (P32352) | steroid biosynthesis |
| YALI1_B29483g | similar to *N. crassa* related to hydroxymethylglutaryl-CoA lyase (Q9P3J2) | synthesis and degradation of ketone bodies |
| YALI1_B30181g | similar to *S. cerevisiae* ERG24 (P32462) | steroid biosynthesis |
| YALI1_D21327g | similar to *S. coelicolor* 4-coumarate:CoA ligase (Q9K3W1) | fatty acid activation |
| YALI1_D22124g | similar to *S. cerevisiae* FAA1 (P30624) | fatty acid biosynthesis |
| YALI1_E13541g | YLIP1 (Q99156) | lipid degradation |
| YALI1_E37118g | similar to *S. cerevisiae* TGL2 (P54857) | lipid degradation |
| YALI1_E37982g | similar to *S. cerevisiae* TGL1 (P34163) | lipid degradation |
| YALI1_E38810g | similar to *S. cerevisiae* DGA1 (Q08650) | TAG synthesis |
| YALI1_E38912g | YIPox1 (O74934) | lipid degradation |
| YALI1_E41315g | similar to *N. crassa* ACL1 (Q8X097) | fatty acid biosynthesis |
| YALI1_F09720g | similar to *S. cerevisiae* (P38137) FAT2 | fatty acid activation |
| YALI1_F38317g | similar to *S. cerevisiae* CEM1 (P39525) | fatty acid biosynthesis |
| **Dimorphism** |  |  |
| YALI1_C31122g | YIOPT1 (Q8WZL3) | oligopeptide transporter |
| YALI1_D28351g | similar to *S. cerevisiae* CTS1 (P29029) | chitinase |
| YALI1_D34785g | similar to *S. cerevisiae* SIN3 (P22579) | histone deacetylase complex component |
| YALI1_E05329g | similar to *S. cerevisiae* SSY5 (P47002) | SPS-sensor system (sensing extracellular amino acids) |

| Miscellaneous |  |  |
| --- | --- | --- |
| YALI1_B21855g | similar to *S. cerevisiae* REG1 (Q00816) | transcription factor (SNF1 pathway) |
| YALI1_D10067g | similar to *C. neoformans* Cts1 (Q8J0A3) | calcineurin temperature suppressor |
| YALI1_E01213g | YIAOX (Q8J0I8) | alternative oxidase, mitochondrial |
| YALI1_F11308g | similar to *S. cerevisiae* HIT1 (P46973) | zinc finger protein |
| YALI1_F26757g | similar to *S. cerevisiae* ASG1 (P40467) | transcription factor |
| YALI1_F38526g | similar to *S. cerevisiae* CCS1 (P40202) | catalase |

\* In strain 1889 no reads could be found that map against YALI1\_B10231g.

**Table S6:**
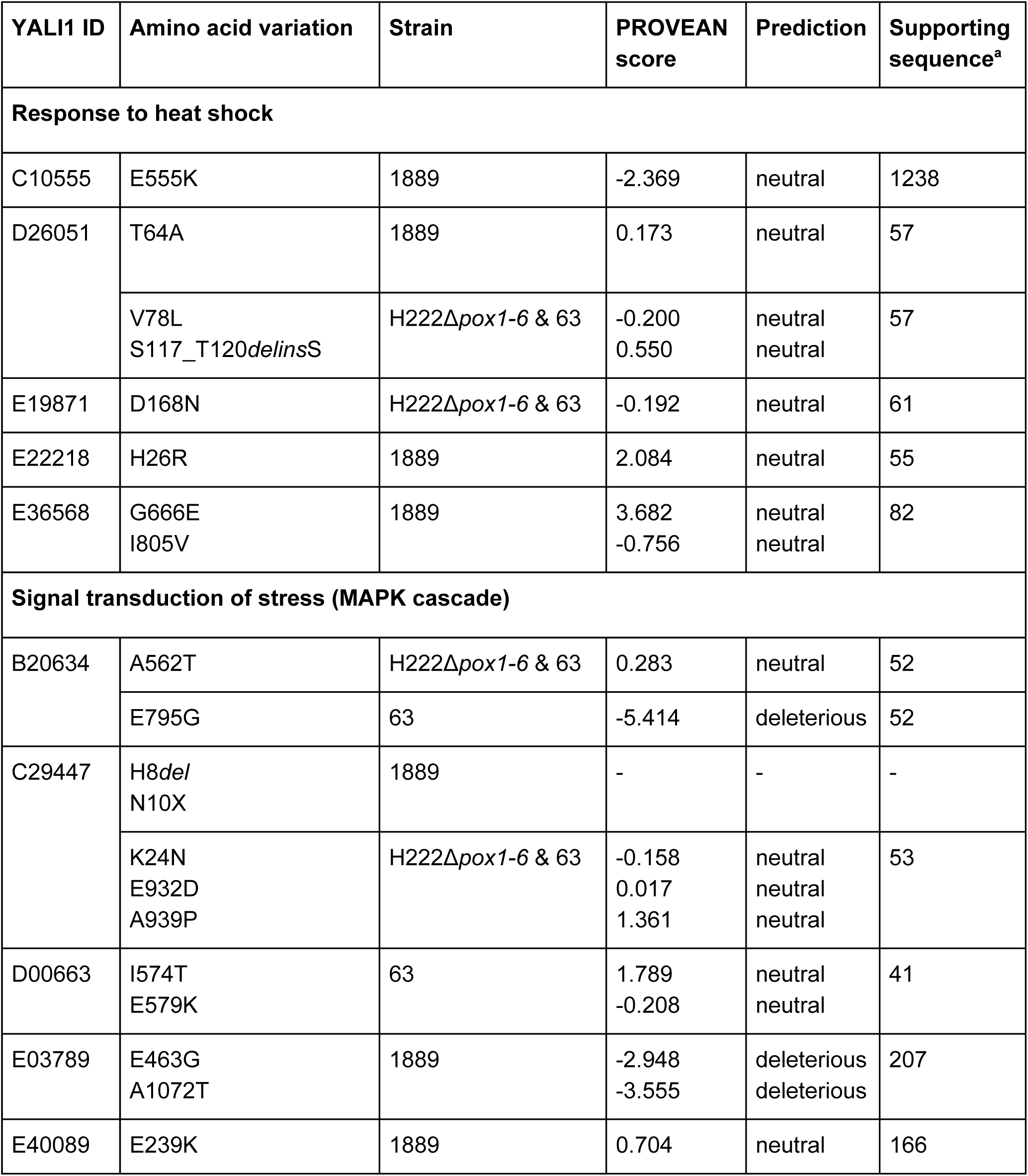
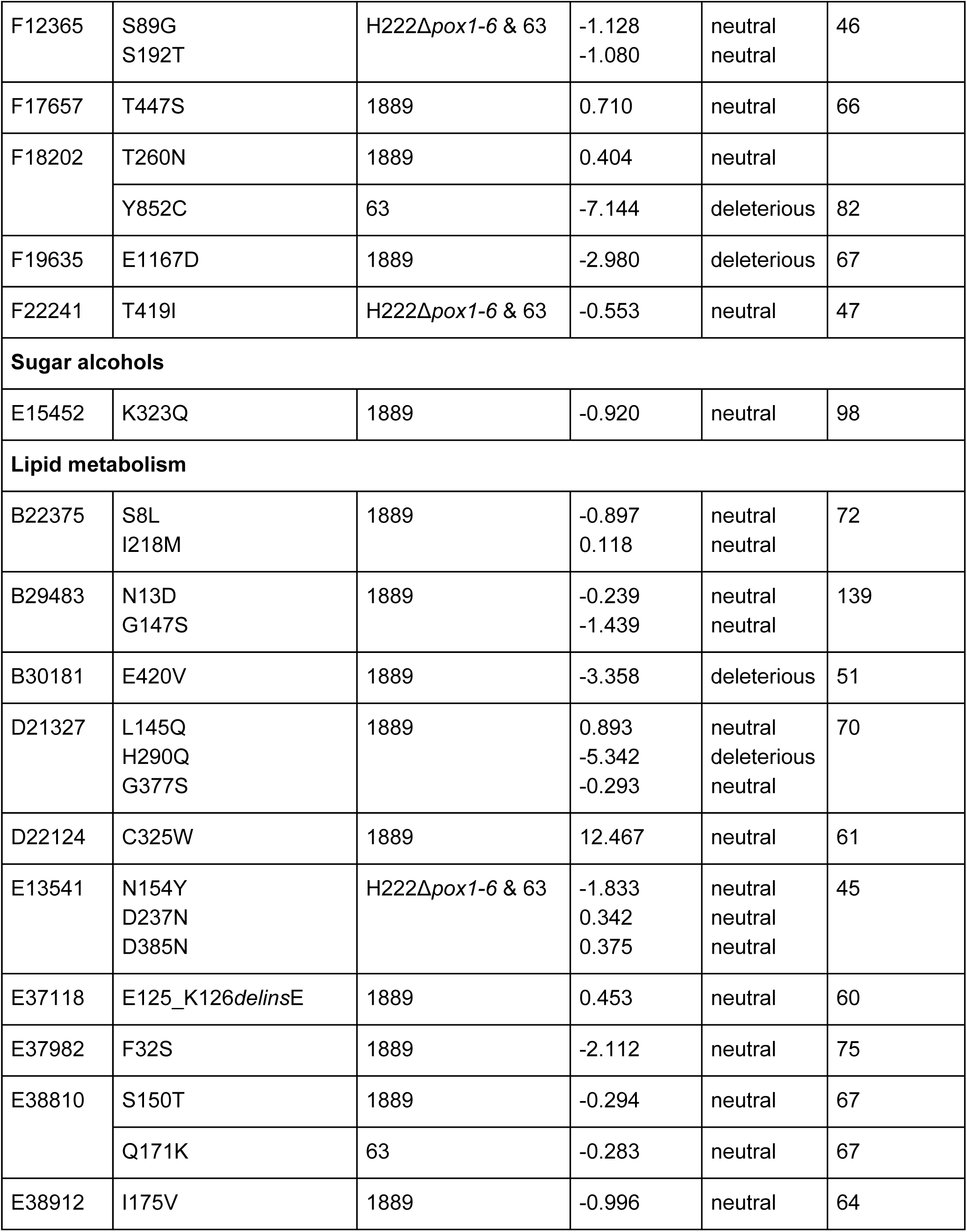
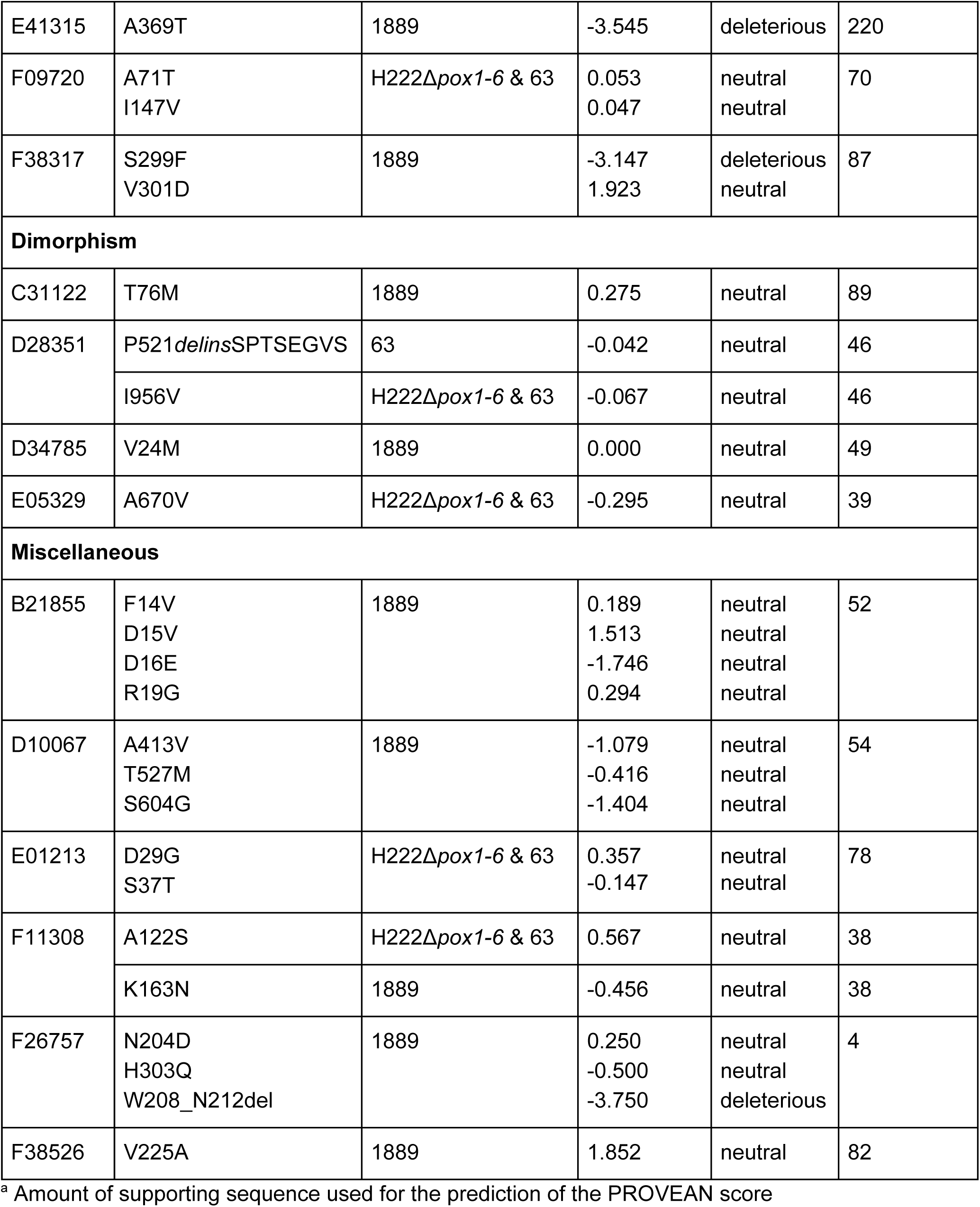
Protein polymorphisms found in *Y. lipolytica* strains H222Δ*pox1-6*, 63 and 1889 and the prediction whether the sequence variations affect the protein functions. Therefore PROVEAN web server was used and the prediction cutoff was set to −2.5 as recommended by the developers. Variants with a score below or equal to the cutoff are considered as “deleterious”.

